# TFEA.ChIP: A tool kit for transcription factor binding site enrichment analysis capitalizing on ChIP-seq datasets

**DOI:** 10.1101/303651

**Authors:** Laura Puente-Santamaria, Luis del Peso

**Affiliations:** Departamento de Bioquímica, Universidad Autonoma de Madrid (UAM) and Instituto de Investigaciones Biomedicas ‘Alberto Sols’ (CSIC-UAM), 28029 Madrid, Spain; IdiPaz, Instituto de Investigacion Sanitaria del Hospital Universitario La Paz, 28029 Madrid, Spain.; CIBER de Enfermedades Respiratorias (CIBERES), Instituto de Salud Carlos III, 28029 Madrid, Spain.

## Abstract

The identification of transcription factors (TFs) responsible for the co-regulation of specific sets of genes is a common problem in transcriptomics. Herein we describe TFEA.ChIP, a tool to estimate and visualize TF enrichment in gene lists representing transcriptional profiles. To generate the gene sets representing TF targets, we gathered ChIP-Seq experiments from the ENCODE Consortium and GEO datasets and used the correlation between Dnase Hypersensitive Sites across cell lines to generate a database linking TFs with the genes they interact with in each ChIP-Seq experiment. In its current state, TFEA.ChIP covers 327 different transcription factors from 1075 ChIP-Seq experiments, with over 150 cell types being represented. TFEA.ChIP accepts gene sets as well as sorted lists differentially expressed genes to compute enrichment scores for each of the datasets in its internal database using an Fisher’s exact association test or a Gene Set Enrichment Analysis. We validated TFEA.ChIP using a wide variety of gene sets representing signatures of genetic and chemical perturbations as input and found that the relevant TF was correctly identified in 103 of a total of 144 analyzed datasets with a median area under the curve (AUC) of 0.86. In depth analysis of an RNAseq dataset, illustrates that the use of ChIP-Seq data instead of PWM-based provides key biological context to interpret the results of the analysis. To facilitate its integration into transcriptome analysis pipelines and allow easy expansion and customization of the TF-gene database, we implemented TFEA.ChIP as an R package that can be downloaded from Bioconductor: https://www.bioconductor.org/packages/devel/bioc/html/TFEA.ChIP.html and github: https://github.com/LauraPS1/TFEA-drafts In addition, make it available to a wide range of researches, we have also developed a web application that runs the package from the server side and enables easy exploratory analysis through interactive graphs: https://www.iib.uam.es/TFEA.ChIP/

## Introduction

Identification of gene expression signatures representing biological states is a key step to understand the transcriptional control of biological processes and the alterations that occur in pathological conditions. The underlying assumption is that one or a few TFs are responsible for the signature. Traditionally, the identification of relevant TFs has relied on the use of position weight matrices (PWMs) to predict transcription factor binding sites (TFBSs) proximal to the DE genes [19]. The comparison of predicted TFBS in DE versus a set of control genes, reveals factors that are significantly enriched in the DE gene set. The prediction of TFBS using these approaches have been useful to narrow down potential binding sites, but can suffer from high rates of false positives. In addition, this approach is limited by design to sequence-specific transcription factors (TF) and thus unable to identify cofactors that bind indirectly to target genes. To overcome these limitations a new family of methods that exploit experimentally determined binding information are beginning to emerge [1] [6]. Here we describe the development the R package TFEA.ChIP, which exploits the vast amount of publicly available ChIP-Seq datasets to determine TFBS proximal to a given set of genes and computes enrichment analysis based on this experimentally-derived rich information. Specifically, TFEA.ChIP uses information derived from the hundreds of ChIP-Seq experiments from the ENCODE Consortium [4] expanded to include additional datasets contributed to GEO database [3] [2] by individual laboratories representing the binding sites of factors not assayed by ENCODE. The package includes a set of tools to customize the ChIP data, perform enrichment analysis and visualize the results. This manuscript describes the main characteristics of the package and assess its performance through the analysis of selected gene sets from the v6.1 MSigDB [10] [9] representing the expression signatures of chemical and genetic perturbations acting through defined transcription factors. Our results show that the relevant transcription factor was identified within the top 10% candidates in 90 out of 129 tested sets. The proportion of correctly identified transcription factors increased up to 13 out of 15 datasets derived from the integration of several signatures. TFEA.ChIP is implemented as a lightweight bioconductor R package facilitating its integration into analysis pipelines and allowing fully customization. In addition, to make it accessible to a wider range of researchers, we have also implemented a web-based tool that runs TFEA.ChIP through a graphic interface.

## Design and implementation

### Database

TFEA.ChIP package includes analysis and visualization tools intended for the identification of TFBS enriched in a set of DE genes. To this end, the package uses information derived from 1075 ChIP-seq datasets, generated by the ENCODE consortium and individual researchers, testing a total of 327 individual human transcription factors in a variety of cell types and experimental conditions (supplemental table S1 Table). The collection of TF includes 276 sequence-specific transcription factors and 51 regulatory molecules that bind DNA indirectly (e.g. transcription cofactors and chromatin modifiers) or in a sequence-independent fashion (e.g. RNA Polymerases). Thus, this compiled database covers around 20% of the 1,391 [17] to 1600 [8] sequence-specific transcription factors encoded by the human genome and include proteins from all the major classes of DNA binding domains (Fig 1 and (supplemental table S2 Table)). Public ChIPseq datasets contain the coordinates of TF binding sites throughout the genome. Thus, in order to use this information in gene enrichment analyses, we first need to associate binding regions (ChIP-peaks) to specific gene loci. In the absence of three-dimensional contact information, such as that produced by chromosome conformation capture carbon copy (Hi-C) experiments, the peaks are usually assigned to the nearest gene.

**Figure 1.**
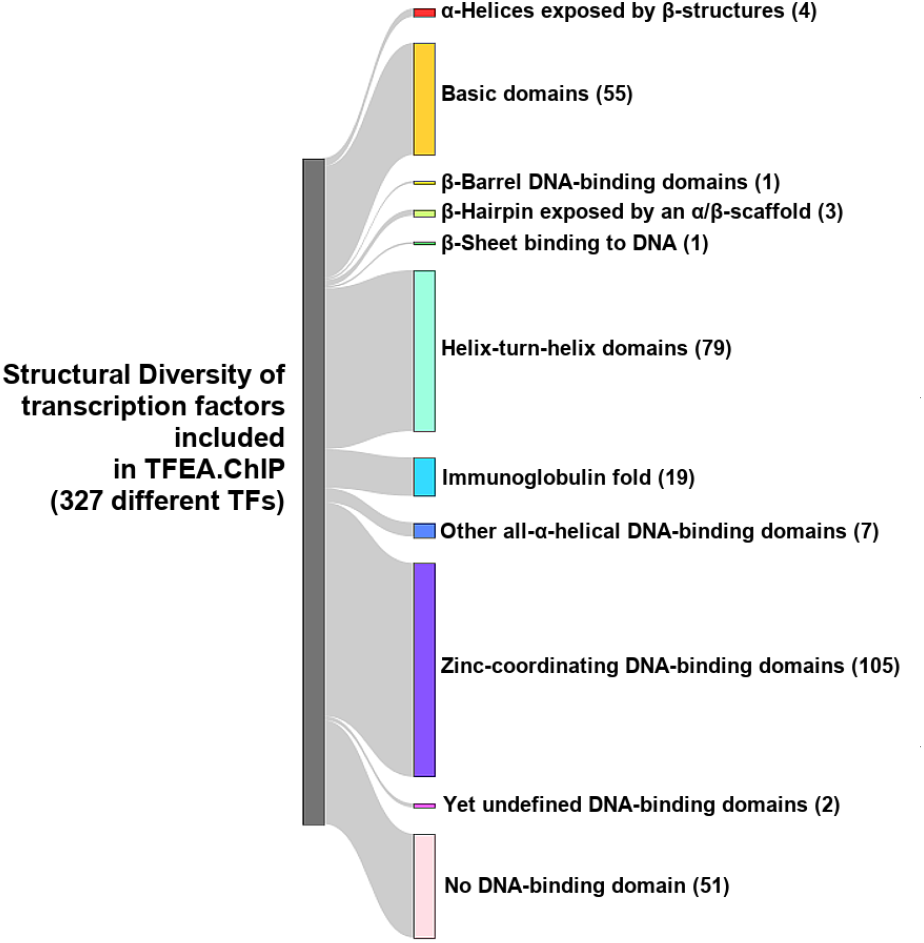
**Structural diversity according to DNA-binding domains of the transcription factors included in the TFBS database.** The 327 TFs included in TFEA.ChIP database were classified into families according to their DNA-binding domain composition.

However, Hi-C experiments indicate that only a small fraction of the looping interactions of distant regulatory regions are with the nearest gene [13]. Accordingly, uncertainty of peak assignation increases as the distanc to the nearest gene broadens [12]. To overcome these difficulties, we exploited the extensive map of enhancer-target gene pairs generated by the ENCODE projec through the analysis of correlation between the DHS signal at distant sites and gene promoters regions across 79 cell lines [15]. Specificall; we first generated a database pairing open chromatin regions, as defined by clusters of Dnase Hypersensitive Sites (DHSs) [15], and genes in the UCSC Known Gene database (version 3.2.2) [5]. DHSs were assigned to genes overlapping with the open chromatin region tolerating a 1Kb margin from the gene boundaries and allowing for multiple gene assignation for those DHSs overlapping two genes. This process resulted in a database of DHSs-gene pairs that only retained DHSs that were assigned to at least one gene (Fig 2, step A). Next, we added to this database the list of statistically significant (Pearson’s correlation coefficient >0.8) enhancer DHSs-gene pairs generated by ENCODE [15] (Fig 2, step B). Then, for each ChlP-seq dataset we selected those peaks that were statistically significant *(FDR <* 0.001 for ENCODE datasets and *FDR <* 0.05 for the rest of datasets) and overlapped an open chromatin region in the DHSs-gene database. Each of these peaks was assigned to the same gene as the DHS they overlapped with (Fig 2, step C). Finally, we integrated the peak-gene information from all ChIP-dataset into a binary matrix with rows corresponding to all the human genes in the Known Gene database, and a columns for every ChIP-Seq experiment analyzed; the entry values were assigned to 1 when the row gene had at least one peak assigned in the ChIP-Seq column and 0 otherwise (Fig 2). It is worth noting that, as a result of the matching strategy, the interaction matrix contains a large fraction of TFBS-gene pairs involving distant regulatory interactions including intronic [12] and enhancer regions.

**Figure 2.**
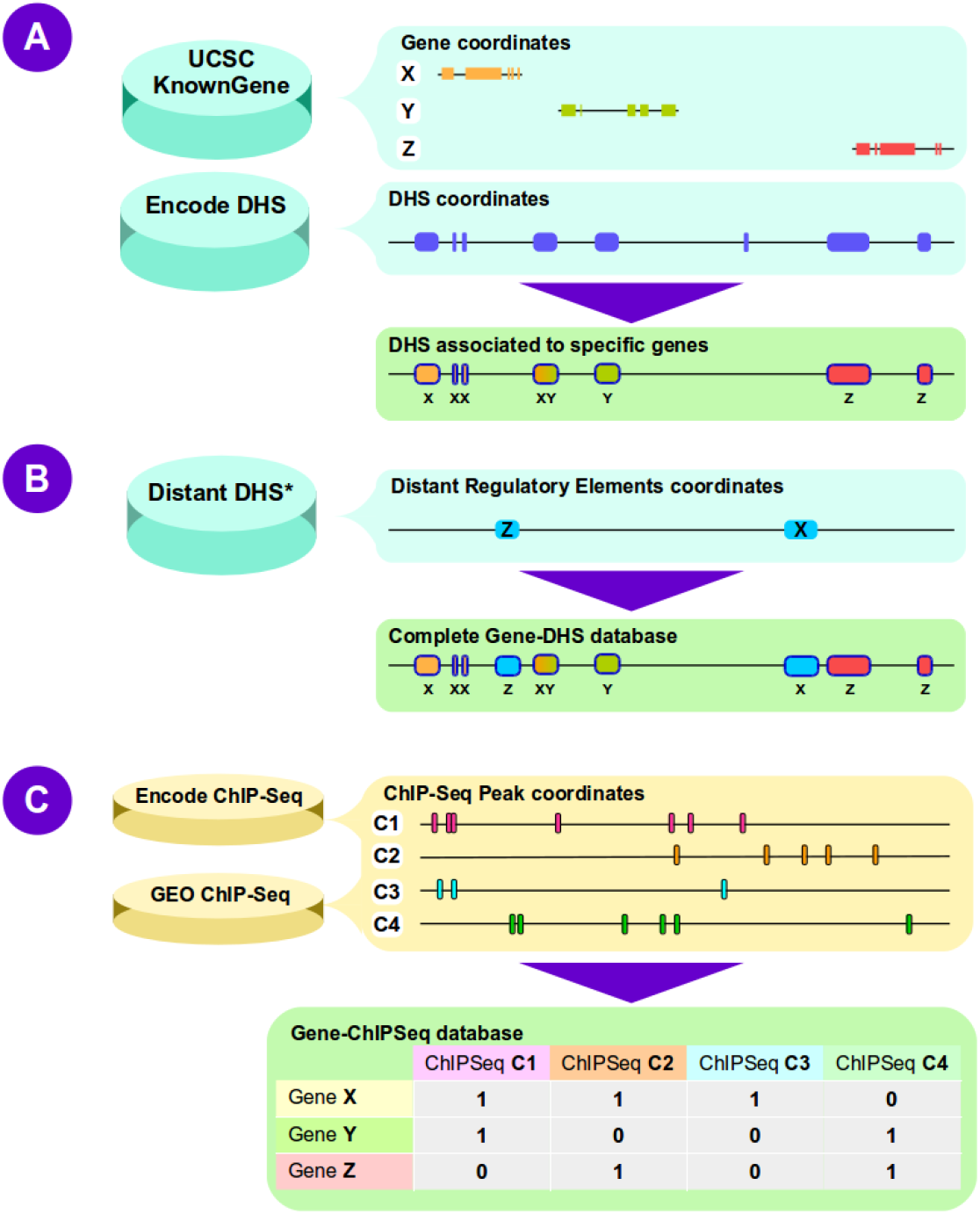
**Building Database of TF-gene associations**. **A**, Open chromatin regions, defined as clusters of DHSs by the ENCODE project, were assigned to the nearest gene in the UCSC Known Gene database within 1Kb window. **B**, Distant DHSs were assigned to genes based on statistical correlation (Pearson’s coeficient >0.8) between distant and promoter DHSs across cell lines. **C**, Significant ChIP-seq peaks mapping to the DHSs selected in A and B were assigned to gene linked to the DHSs they overlap with.

### Enrichment analysis

TFEA.ChIP is designed to take the output of gene expression profiling analysis and identify transcription factors enriched in the list of differentially expressed genes. The core premise of the method is that key effectors of a transcriptional response will have more target genes among the differentially expressed than among the unresponsive genes. TFEA.ChIP implements to types of tests to identify enriched TF. The first one analyzes the association of TFBS and differential expression from 2x2 contingency tables categorizing all human genes according to the presence of binding sites for a given TF and their transcriptional response (DE or non-responsive). The statistical significance of the association for each transcription factor is determined by a Fisher’s exact test. We refer to this method as *association test* throughout the text. This analysis only requires a list of DE genes as input and returns a table containing the results of the Fisher’s exact test computed for each one of the 1075 independent binding profiles the data base. In addition to the FDR-adjusted p-value (and its −log10 transformation, here referred to as LPV) for the association, the table contains the odds-ratio (and the log2 transformation, here referred to as LOR) as a measure of the size effect. Also, in an attempt to rank the transcription factors as potential candidates mediating the regulation of DE genes, the program computes the euclidean distance of each factor i to the origin in LPV versus LOG graphs as

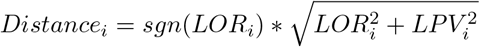

where *LOR_i_* and *LPV_i_* correspond to the log_2_*(Oñ)* and — (log_10_ (*adjPval*)) transformation of the odds-ratio (OR) and FDR-adjusted p-value (adjPval) returned by the Fisher’s exact test for the *i* TF in the data base, and *sng()* is the sign function. In the second method, the association of TF to DE genes is determined using a Gene Set Enrichment Analysis (GSEA) [14]. To this end TFEA.ChIP uses the list of genes bound by each TF in the internal database as a gene sets representing the binding signature of each factor. Thus, each column in figure 2 is represented as a gene set that includes all genes with a value of 1. This analysis requires as input list of genes sorted according to the magnitude of the difference in expression in the two conditions being compared. We recommend the π-value [20], which combines expression fold change and statistical significance, as sorting criteria for the result of a DE gene analysis:

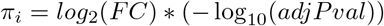

where FC is the ratio of gene *i* expression in the two conditions being compared and adjPval is the multiple-testing adjusted P-value associated with equal-mean expression hypothesis test.

## Results

The general strategy followed by TFEA.ChIP was first described in an study aiming to determine transcription factors involved in gene repression induced by reduced oxygen availability (hypoxia) [16]. This work also highlighted the benefits of using ChIP-seq data over PWM for the identification of TFBS [16]. The implementation of this approach in the package TFEA.Chip described herein greatly simplifies its application to any general case. Here, we use a case study to illustrate the tools implemented in this software package and its web version. In addition, to provide evidence of its general performance, we applied TFEA.ChIP to a total of 144 gene sets from the MSigDB representing the expression signature of a wide variety of chemical and genetic perturbations and well-defined lists of transcription factor targets.

### 0.1 Case study: identification of TF responsible for hypoxia-triggered gene induction

The transcriptional response to hypoxia is mediated by a group of basic helix-loop-helix (bHLH) transcription factors termed Hypoxia Inducible Factors (HIFs). HIFs are heterodimers that share a common beta subunit, encoded by the gene *ARNT*, and an alpha subunit, encoded by the genes *HIF1A, EPAS1* or *HIF3A*. To demonstrate the use of TFEA.ChIP we applied it to an RNA-seq dataset (GSE89831) representing the transcriptional response of endothelial cells to hypoxia [16]. To this end, we first reanalyzed this dataset with DEseq2 [11] and selected DE genes whose transcription was significantly induced by hypoxia (log-fold change hypoxia over normoxia >0 and FDR <0.05). Then, we searched for overrepresented TFBS in this list of DE genes using the association test in TFEA.ChIP (Fig 3A) and Opossum (Fig 3B), a PWM-based state-of-the-art tool [7]. To compare both methods, we processed the raw output produced by oPOSSUM (number of target hits, target non-hits, background hits, and background non-hits for every PWM) to generate statistics comparable to those produced by TFEA.ChIP (Fisher’s tests p-value and odds ratio). While both methods were able to find the Hypoxia Inducible Factors (HIFs) as TF significantly enriched in the set of genes upregulated by hypoxia (Fig 3A and B), the results of TFEA.ChIP clearly ranked HIFs above the rest of TFs, suggesting a much higher signal to noise ratio. In addition, the output of TFEA.ChIP indicated that most of the datasets representing these transcription factors are consistently enriched (Fig 3A), regardless the HIF subunit assayed (HIF1A, EPAS1 and ARNT, no ChIP-seq datasets are available for HIF3A). In fact, the interactive analysis allowed by the package, revealed that the datasets that did not show significant enrichment correspond to samples assayed in normoxic conditions where HIFs are inactive (Fig 3C). Conversely, the few datasets from normoxic samples that showed enrichment derive from a clear cell renal carcinoma cell line (786-O). This cell line is defective for the tumor supressor gene VHL, a key protein controlling the transcriptional response to hypoxia, leading to constitutive activation of HIF even in the presence of oxygen (normoxic conditions). The association analysis implemented in TFEA.ChIP allows ranking the ChIP datasets based on their distance to the origin in the LPV-LOR plots (see mehods). Based on this information, we found that, in agreement with the observations above, HIF (ARNT, EPAS1 and HIF1A) datasets derived form hypoxic experiments significantly ranked above other TF datasets (*p — value* = 1.6e — 6, Mann-Whitney U test). In addition to the association analysis, TFEA.ChIP includes a GSEA-based analysis to determine the enrichment of TFBS on a sorted list of genes. To apply this analysis to the hypoxia gene expression data set, we first sorted the genes detected in the experiment according to their response to hypoxia (pi-value, [20]) and then used the sorted list as input for the GSEA included in the package. Figure 4A shows the GSEA-derived enrichment score (ES) and rank for the ES (Argument) for each one of the 1075 gene sets representing the signatures for each one of the TF included in the database. The highest scoring significant gene sets correspond to those representing HIF binding profiles (figure 4A). Accordingly, the distribution of genes bound by HIF is strongly skewed toward large values of *π* in the sorted DE gene list 4B).

**Figure 3.**
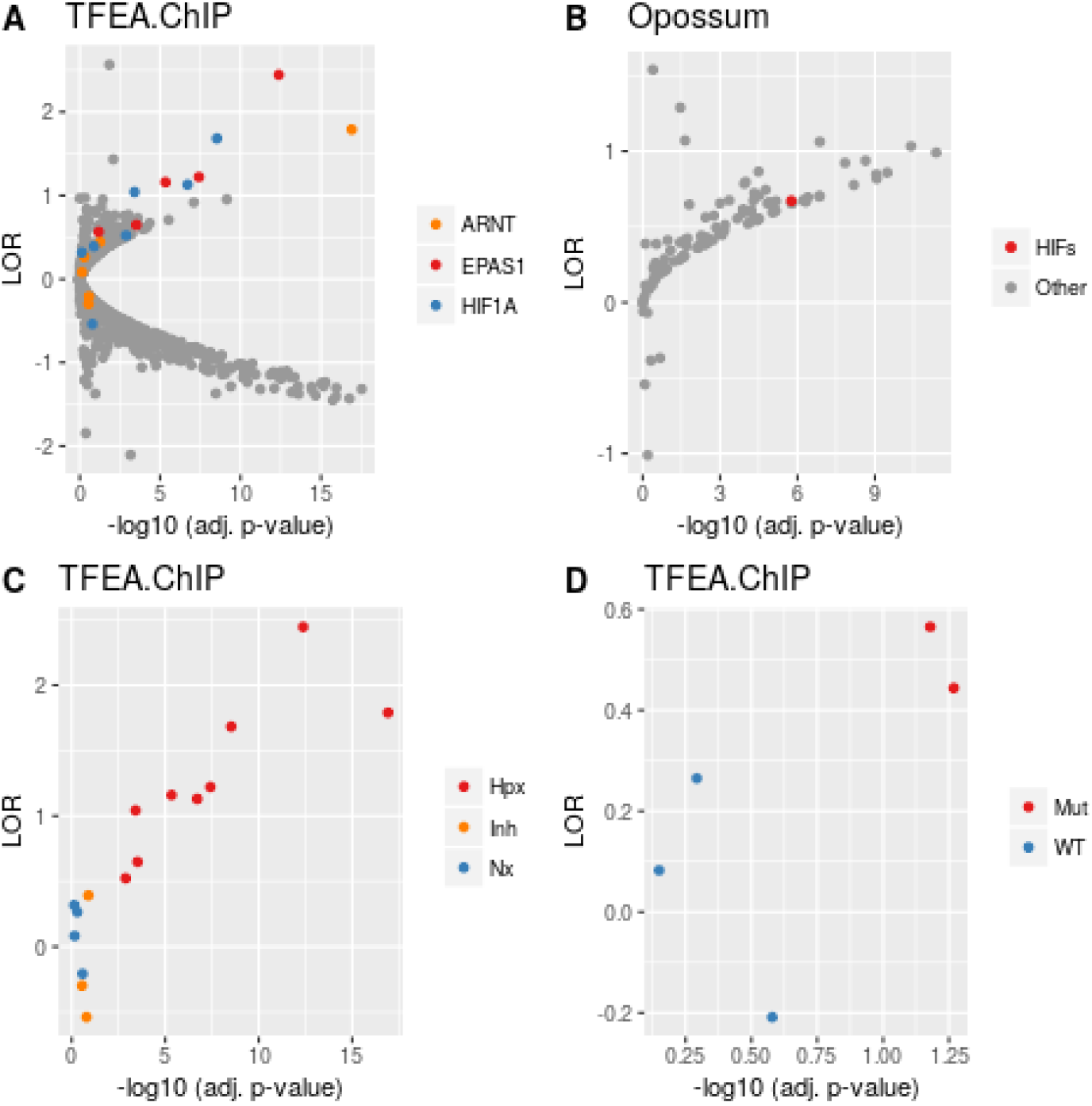
**Identification of TF enriched in genes induced by hypoxia. Association test.** A set of genes whose transcription was significantly induced in response to hypoxia was analyzed with TFEA.ChIP (A,C and D) or Opossum (B). The graphs represent the adjusted p-value (-log10 FDR) and the log-odds ratio (LOR) for the association of ChIP datasets (A,C and D) or PWM-motifs (B). **A**, datasets corresponding to TF of the Hypoxia Inducible Factor (”HIF1A”, “EPAS1” and “ARNT”) family and other TF (”other”); **B**, PWM-motif corresponding to Hypoxia Inducible Factors (”HIFs”) vs rest of motifs (”Other”). **C**, the graph shows HIF datasets labelling those derived from normoxic (”Nx”), hypoxic (”Hpx”) and inhibitor-treated (”Inh”) samples. Inh samples were exposed to a small molecule inhibitor that activates HIF. **D**, HIF datasets labelling those derived from VHL-competent (”WT”) or deficient (”Mut”) cells.

**Figure 4.**
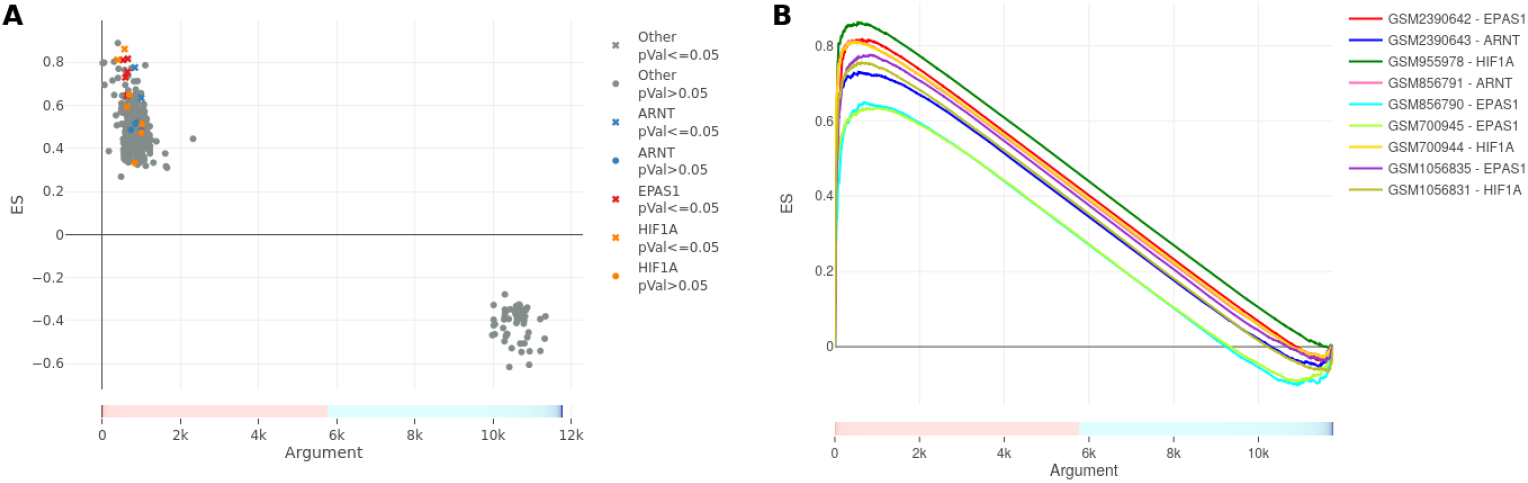
**Identification of the TF responsible for transcriptional upregulation induced by hypoxia. GSEA analysis**. We computed the π-value [20] for the genes in the DE analysis and used it as the sorting variable in the GSEA analysis. **A**, the graph represents the ES for the 1075 TFBS included in TFEA tested against the list of genes sorted by their response to hypoxia. Gene sets with ES values that differ significantly from random associations are shown as crosses and non-significant values as circles. Gene sets representing the binding of HIFs are shown in color (EPAS1, red; HIF1A, orange; ARNT, blue) and the rest of factors in grey. **B**, The graph represents the profile of the running ES for the indicated gene sets representing HIFs.

Altogether these results indicate that the analyses implemented in TFEA successfully identified HIFs as the relevant factors driving the transcriptional upregulation induced by hypoxia and show that the biological metadata associated to the ChIP datasets included in the package provide invaluable context to interpret the output.

### 0.2 General performance of TFEA.ChIP

To determine the general performance of TFEA.ChIP, we tested it against the annotated gene sets in the Molecular Signature Database (MSigDB) [14] [10]. Specifically, we selected 129 gene sets included in the C2-CGP (”curated datasets”) collection, representing expression profiles of defined genetic and chemical perturbations, and used them as input for TFEA.ChIP. The selection criteria included gene sets from experiments that targeted a defined TF and that the factor (or a paralog) was present in the internal database of TFEA.ChIP. The selected gene sets include a variety of experimental settings affecting 34 families of TF (Supplementary table S3 Table). To visualize the performace of TFEA.ChIP on these datasets we recorded the rank occupied by the relevant TF in the ouput of the association analysis. As shown in Fig 5A (”C2” group), the relevant TF was present within the top 10% ranking candidates for the majority (90 out of 129) of analyzed datasets. The C2 subset of the MSigDB derive from individual experiments performed by independent researchers using a wide array of techniques and settings and hence it is a heterogeneous and redundant collection. Thus, we next tested TFEA.ChIP against the MSigDB “Hallmark” collection that consists of a non-redundant and manually curated gene sets representing specific biological states [9]. In this case, the relevant TF was present in the top 10% ranking factors in 13 out of 15 gene sets representing defined gene expression signatures (Fig 5A “H” group). In addition, the relevant TFs were usually found far from the remaining TF in the LPV-LOG plot (Fig 5B), suggesting a good signal-to-noise ratio. In agreement, computation of receiver operating characteristic (ROC) curve for each dataset showed that the area under the curve (AUC) was higher than 0.7 for 73% of the C2 and 100% of the Hallmark datasets (Fig 5C). In the case of the Hallmark collection, the AUC was >0.8 for 14 of the 15 gene sets and >0.9 for 7 of them. Thus, TFEA.ChIP shows a high discriminative capacity across a wide range of gene profiles derived from varied experimental conditions and cell origins.

**Figure 5.**
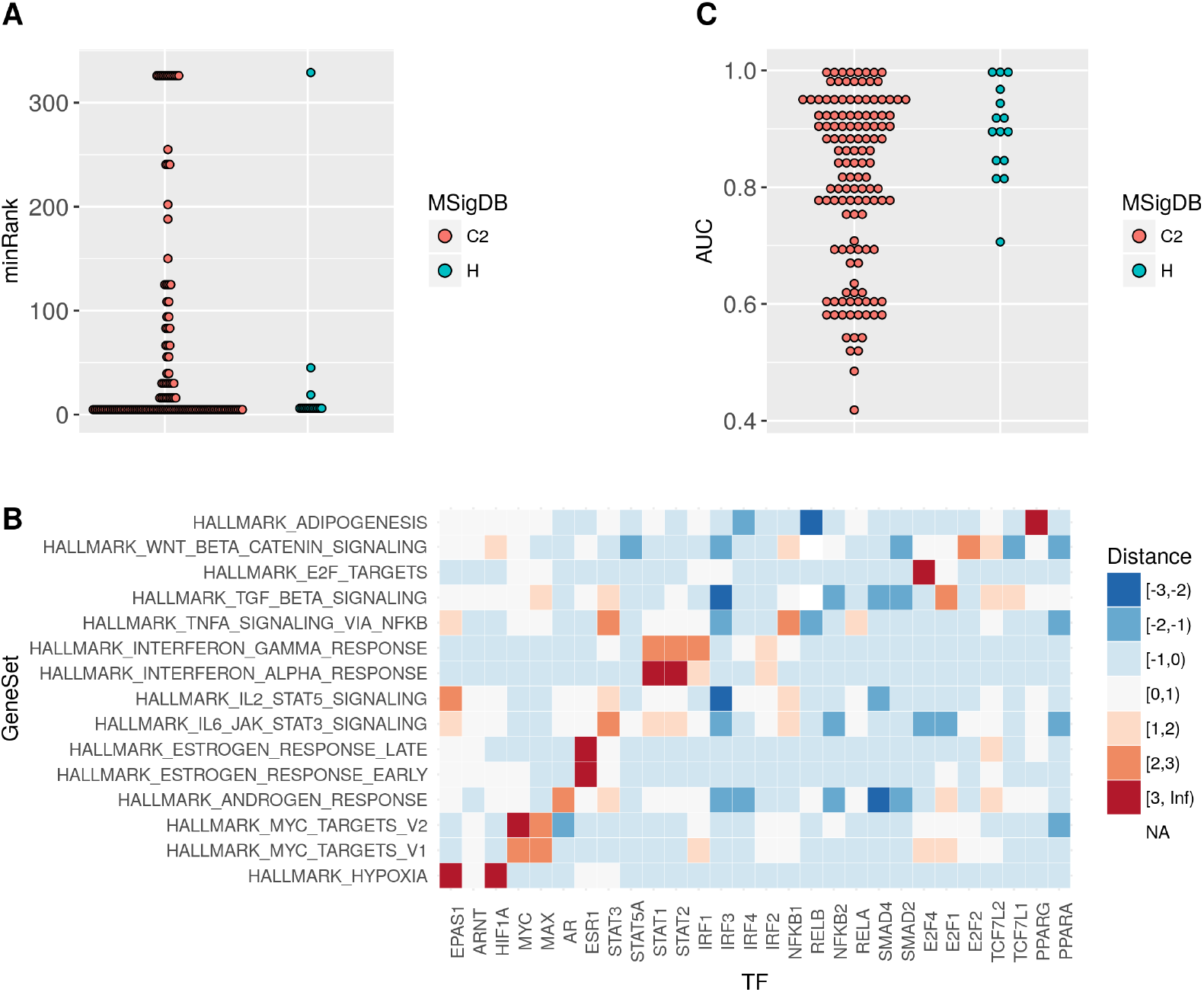
**Performance of TEFA.Chip on MSigDB gene sets**. **A**, Selected gene sets from the C2 and H collections of the MSigDB were used as input for the association test. The distance of each ChIP data set to the origin was computed as indicated in methods and the rank of the relevant TF in the sorted list of distances was recorded. The graph represent the minimum rank among datasets representing the relevant factor. The indicated gene sets from the Hallmark collection were used as input for the association test of TFEA.ChIP. **B**, For each analysis, the distances to origin of all ChIP datasets were normalized to standard scores. The heatmap represents the maximum standard score among the ChIP dataset representing the indicated TF across the tested gene sets. **C**, For each tested gene set, we computed a ROC from the sorted list of distances of each ChIP dataset set to the origin, labeling as true positives those datasets corresponding to the relevant TFs. The graph represent the AUC for analyzed gene sets.

## Discussion

The identification of the transcription factor(s) that coordinate a given gene expression pattern is usually a key piece of information in transcriptomics. Herein, we describe the package TFEA.ChIP, a software package that combines experimentally determined genome-wide binding profiles of transcription factors and DNase hypersensitive regions to identify TFBS enrichment. The use of ChIP-seq data instead of binding profiles in the form of PWM, to determine gene-regulatory networks is an emerging approach and, to our knowledge, just two other tools, ENCODE ChIP-seq significance tool [1] and iREGULON [6], make use of it. However, both packages significantly differ from TFEA.ChIP in the strategy they use to identify enriched TFBS and, importantly, their implementation. A main difference is that TFEA.ChIP makes use of correlation between DHSs across different cell types to assign TFBS to genes, instead of just assigning the ChIP-peaks to the nearest gene/feature within a defined window. Another important difference is that the database containing TFBS-genes pairs that TFEA.ChIP uses to compute associations and enrichment scores is fully customizable. The software package includes functions to allow users to expand the internal database and incorporate additional criteria, such as histone postranslational modifications, to assign ChIPseq peaks to genes. Finally, ENCODE ChIP-seq significance tool and iREGULON are implemented as web applications (in the case os iREGULON as a Cytoscape plugin that connects to a server-side daemon over the Internet). In contrast, TFEA.ChIP is a lightweight R package that includes data base and analysis tools, allowing its integration with other libraries in general transcriptomics pipelines. In addition, to make it available to a wide range of researchers, we used Shiny (https://shiny.rstudio.com/ to built an interactive web application that enables the use of TFEA.ChIP analysis tools through a simple graphic interface. The user guide for the web application, including an step-by-step analysis of the hypoxia dataset is presented in the supplemental information S1 File. A limitation of the current version of the TFEA.ChIP package is that it only includes ChIP datasets derived from human cells. To circumvent this restriction, the package includes a function to translate mouse gene names to their equivalent ID on the human genome, enabling the analysis mouse datasets. Since many of the genes representing particular signatures are conserved between human and mouse, translation of terms should not affect the ability to identify the relevant TF, as indicated by our preliminary analysis (data not shown). In conclusion, TFEA.ChIP is an R package that exploits experimentally determined genome-wide binding profiles to accurately predict the TF(s) that mediate gene signatures or transcriptional profiles. During the preparation of this manuscript Wang et al. published a manuscript describing another software tool, BART, that leverages on publicly available ChIP-seq profiles to predict transcription factor enrichment [18]

## Supporting Information

### S1 Table

**ChIP-seq datasets included in the TFEA.ChIP package**. The table includes the datasets along with their GEO accession ID and metadata.

### S2 Table

**Structural classification of the transcription factors included in TFEA.ChIP**.

### S3 Table

**Gene sets from MSigDB analyzed used to test TFEA.ChIP performance**. The table includes MSigDB gene set ID (“Gene_Set”), reference to the collection name (“Collection”), the genetic/chemical perturbation or biological/experimental process represented by the gene set (“Treatment”) and the main transcription factor(s) expected to be activated by the perturbation as well as paralogs and related factors (“TF”).

### S1 File

**TFEA.ChIP web application user guide**.

## Acknowledgments

We thank Javier Merino for his help with the web application. This work was supported by Ministerio de Ciencia e Innovation (Spanish Ministry of Science and Innovation, MICINN) [SAF2014-53819-R to LdelP, SAF2017-88771-R to LdelP]; by Fundacion Caja Madrid (Beca de Movilidad para Profesores de las Universidades Publicas de Madrid 2011–2012 to LPO).

